# G-quadruplex dynamics contribute to epigenetic regulation of mitochondrial function

**DOI:** 10.1101/359703

**Authors:** M. Falabella, J. E. Kolesar, I. M. Xiang, T. Wang, W. Horne, C. Wallace, L. Sun, Y. V. Taguchi, C. Wang, J. Turek-Herman, C. M. St. Croix, N. Sondheimer, L. A. Yatsunyk, F. B. Johnson, B. A. Kaufman

## Abstract

Single-stranded DNA or RNA sequences rich in guanine (G) can adopt non-canonical structures known as G-quadruplexes (G4). Predicted G4-forming sequences in the mitochondrial genome are enriched on the heavy-strand and have been associated with formation of deletion breakpoints that cause mitochondrial disorders. However, the functional roles of G4 structures in regulating mitochondrial respiration in non-cancerous cells remain unclear. Here, we demonstrate that RHPS4, previously thought to be a nuclear G4-ligand, localizes primarily to mitochondria in live cells by mechanisms involving mitochondrial membrane potential. We find that RHPS4 exposure causes an acute inhibition of mitochondrial transcript elongation, leading to respiratory complex depletion. At higher ligand doses, RHPS4 causes mitochondrial DNA (mtDNA) replication pausing and genome depletion. Using these different levels of RHPS4 exposure, we describe discrete nuclear gene expression responses associated with mitochondrial transcription inhibition or with mtDNA depletion. Importantly, a mtDNA variant with increased anti-parallel G4-forming characteristic shows a stronger respiratory defect in response to RHPS4, supporting the conclusion that mitochondrial sensitivity to RHPS4 is G4-structure mediated. Thus, we demonstrate a direct role for G4 perturbation in mitochondrial genome replication, transcription processivity, and respiratory function in normal cells and describe the first molecule that differentially recognizes G4 structures in mtDNA.

## Introduction

Mitochondria are essential, flexible organelles that play a crucial metabolic role in stress and disease resilience. Mitochondria are also bigenomic, with subunits of the large multimeric oxidative phosphorylation (OXPHOS) complexes encoded by both nuclear and mitochondrial DNA (mtDNA). Genetic lesions arising in either the Mendelian nuclear genome or the matrilineal multicopy mtDNA can result in mitochondrial dysfunction. Mammalian mtDNA is a circular genome characterized by strand asymmetry with guanine enrichment in the “heavy” strand, which has implications for genome stability. During first strand mtDNA replication, the heavy strand is displaced from the protective protein scaffold normally associated with mtDNA (1), increasing the potential for its damage. Importantly, the primary driver of age-acquired mutation is thought to be cytosine deamination, whereas guanine oxidation is not a major mutational event in mtDNA (2, 3). The heavy-strand guanine enrichment may thus reduce the incidence of cytosine to thymidine transitions on this strand (4, 5).

One potential effect of guanine enrichment in the heavy strand may be the increased potential for intramolecular G-quadruplex (G4) formation during replication. G-quadruplexes are predicted when four runs of two or more consecutive guanines occur nearby one another within a nucleic acid sequence. The frequency of sequences with such G4-forming potential in mtDNA appears to be non-random (6), with many potential G4s being conserved and verified to form G4 structures *in vitro* (6, 7). During first strand mtDNA replication, the heavy strand is displaced, providing an opportunity for the assembly of G4 structures. Second strand replication utilizes the displaced heavy strand as template, creating the potential for replication pausing through template interference. Toward understanding the origin of mtDNA deletions, we and others have described G4-forming sequences associated with mtDNA deletion breaks points from patients with mitochondrial disorders (6–8).

The current model of G4 structure involvement in mtDNA instability, specifically in the formation of deletions, is largely inferred from observations of their effect on nuclear DNA. For example, small molecule G4-ligands that bind and stabilize G4 structures can induce genomic instability at sequences with high potential for G4 formation, such as the human minisatellite CEB1 (9) and telomeric DNA (10, 11). Cells possess ATP-dependent helicases that are able to resolve these G4 structures (12). The Pif1-family of G4 helicases was first identified in budding and fission yeast, and localizes to both the nuclear and mitochondrial genomes (13, 14) and binds and resolves G4 structures to prevent replication pausing (9, 15) and chromosome sequence instability (16). In humans, PIF1 acts within the normal nuclear genome, at minisatellite sequences, and at telomeres (17, 18). In cancer cells, there is a strong association between the positions of nuclear genome deletion breakpoints and sequences predicted to form G4 structures, suggesting that replication pausing is a contributing factor in deletion formation (19).

The association of predicted G4-forming sequences with mtDNA deletions does not prove that G4 structures cause nucleic acid instability *per se*, nor does it define the potential roles G4s have on biological function. A mitochondrial G4 biological effect was suggested when we and our collaborators described a precocious yet subtle increase in mtDNA deletion burden in the skeletal muscle of PIF1 G4-helicase ablated mice (20), in which both the nuclear and mitochondrial PIF1 isoforms are eliminated (21). However, the small increase in mtDNA deletions does not explain the notable respiratory complex deficiency in skeletal muscle of those mice (20) or in PIF1 knockdown cells (22). Intriguingly, data from other groups show that in *in vitro* mitochondrial transcription assays, hybrid G4 structures forming in the nascent transcript can cause transcript termination, which suggests that G4 formation could alter gene expression (23–25). In mtDNA, there are many non-randomly distributed and well-conserved sequences with G4-forming potential, which compelled further investigation into the potential biological impact of G4 structures on mitochondrial function.

Despite these connections, the role of G4 in mtDNA is unclear, requiring the development of novel reagents for their study (26). As such, we sought a means to alter mitochondrial G-quadruplex stability and assess the effect of such an alteration on mtDNA metabolism and respiratory function in normal cells. Recently, the small molecule BMVC, possessing *in vitro* G4 binding activity, has been shown to localize to mitochondria and cause apoptosis in cancer cell lines, but not in normal cell lines (27). Here, we here demonstrate that a different compound, RHPS4, previously considered as a nuclear G4 ligand, is preferentially localized to mitochondria where it functions as a G4-ligand in both cancerous and non-cancer cell lines. RHPS4 localization to mitochondria involves both membrane potential and retention by mitochondrial nucleic acids. We describe the ability of RHPS4 to increase antiparallel G4 structure stability of mtDNA and differentiate between genome variants with based on G4-forming potential to affect mitochondrial genome content, transcription elongation, and respiratory function. Leveraging the ability of RHPS4 to cause transcription or replication defects depending on concentration, we identify novel gene expression pathways that distinguish between these processes. Together, these results implicate mitochondrial G-quadruplex stabilization as a negative regulator of mitochondrial nucleic acid synthesis and mitochondrial function.

## MATERIALS AND METHODS

### Chemicals

<u>Cell culture reagents</u>: Dulbecco’s Modified Eagle’s Medium (DMEM; HyClone, Logan, UT); trypsin-EDTA 1X 0.25% (Corning, Tewksbury, MA); bovine serum albumin (BSA) and ethidium bromide (EtBr; Sigma-Aldrich, St. Louis, MO); fetal bovine serum (FBS); fetal calf serum (FCS); phosphate buffered saline (PBS) and penicillin/streptomycin (ThermoFisher Scientific, Waltham, MA). <u>Chemicals</u>: 3,11-Difluoro-6,8,13-trimethylquino[4,3,2-*kl*]acridinium methylsulfate (RHPS4; kind gift of Dr. Marc Hummersone, Pharminox Ltd., Nottingham, UK and purchase from Tocris); BRACO-19 (Organic Synthesis Facility at Fox Chase, Philadelphia, PA); N-methyl mesoporphyrin IX (NMM; Frontier Scientific, Logan, UT); Phen-DC3 (kindly provided by Dr. David Monchaud and Dr. Marie-Paule Teulade-Fichou, Curie Institute, Paris, France); FCCP (trifluoromethoxy carbonylcyanide phenlhydrazone; SigmaAldrich, St. Louis, MO); and cyclosporin A (CsA; Fisher Scientific, Pittsburgh PA). All primers and probes are from Integrated DNA Technologies (Coralville, IA). All other chemicals were reagent grade or better.

### Cell culture and treatments

Mouse embryonic fibroblasts (MEFs), C2C12 myoblasts, HeLa, 143B ρ^0^ and 143B ρ^+^ cells were grown in DMEM medium supplemented with 25 mM glucose, 4 mM glutamine, 110 mg/ml pyruvate, 10% (v/v) FBS:FCS (1:1), 100 U/ml penicillin and 100 mg/ml streptomycin at 37°C under standard conditions (5% CO_2_; ambient O_2_; 95% relative humidity). 143B ρ^0^ and 143B ρ^+^ medium was further supplemented with 0.2 mM uridine. HeLa cells were verified by single nucleotide polymorphism genotyping at the University of Pittsburgh HSCRF Genomics Research Core. All cell lines were mycoplasma free. Sub-confluent wild type MEFs and HeLa cells were cultivated in media containing RHPS4 at indicated concentration (0, 1, 2, or 10 μM) and time (16 or 24h). Drug wash-out was performed by pre-warmed media replacement. EtBr treatment was conducted on sub-confluent MEFs for 24 h in the presence of 25 or 250 ng/ml compound. Cells were collected by trypsinization, pelleted at 200g for 5 min, washed with PBS, and flash frozen in liquid nitrogen for later DNA or RNA preparation.

### Cell viability measurement

Cell viability was assessed by crystal violet assay as reported in (28). Briefly, 143B ρ^+^ and ρ^0^ cells were seeded in 96 wells plate at 5 × 10^3^ cells/well, eight wells per group. After 24 hrs incubation with RHPS4 (0; 2;10 μM) or CsA (0; 0.5; 1; 2;10 μM), the medium was discarded, the cells washed with PBS and incubated with a crystal violet staining solution (0.5%) at room temperature for 20 min. The cells were then washed twice with water and dried for 2 hrs. Dye was released by 20 min incubation in 200 μl methanol and the optical density of stained cells was read at 570 nm using a BioTek Sinergy 4 Hybrid Multi-Mode Microplate Reader (BioTek, Winooski, VT; USA).

### Identification of mitochondrial sequences with G-quadruplex forming potential

Reference mouse mtDNA sequence (NC_005089) was subjected to G4 Hunter QFP prediction as using 25 nt window and a threshold of 1.0 (29). Merged outputs are shown with any overlapping results adjusted to be discrete.

### mtDNA measurements

Total genomic/mitochondrial DNA was isolated by sodium dodecyl sulfate lysis and proteinase K digestion from MEFs and C2C12 as previously described (30). DNA was gently resuspended at 37°C in Tris-EDTA (TE) buffer supplemented with RNAse A to remove the RNA contamination and DNA concentration was measured using AccuBlue Broad range kit (Biotium, Fremont, CA). For measuring relative mouse mtDNA abundance, we designed TAQMAN primer/probes for mitochondrial ND1 (VIC-labeled; primer:probe of 1:1)) and nuclear TBP (FAM-labeled; primer:probe of 3:1), screened candidates for *in silico* primers compatibility, and validated the multiplex reaction by template serial dilution. The multiplex assessment of mtDNA relative abundance content was conducted by qPCR using TaqMan Fast Advanced Master Mix (ThermoFisher Scientific), 4.6 ng/reaction DNA and 5 μM of primer/probes in 10 μl final reaction volume and calculated by the ΔΔC_q_ method (31). The qPCR amplification profile was: one cycle (95°C for 20 sec) and 40 cycles (95°C for 1 sec and 60°C for 20 sec).

Measurement of major mitochondrial topoisomers was performed as previously reported (30, 32). In brief, total DNA was isolated as above, quantified by Nanodrop 1000 (ThermoFisher Scientific), and 2 μg of DNA resolved on 0.7% gel in 0.5X Tris/Borate/EDTA (TBE) buffer for 17 hours at 40V. DNA in gel was nicked, denatured, neutralized, and capillary transferred onto positively charged nylon (Hybond N+; GE Healthcare, Marlborough, MA). Membrane was rinsed, UV crosslinked, and air dried. MtDNA isoforms were detected after random-hexamer probing and quantifying bands by phosphorimager analysis.

For the *in organello* replication assay, mitochondria were isolated from WT mouse liver by differential centrifugation (33) and protein determined by Qubit fluorescent assay (ThermoFisher Scientific). Mitochondria were pelleted and resuspended at 4 mg/ml protein concentration in incubation buffer (10 mM Tris-HCl pH 8.0, 20 mM sucrose, 20 mM glucose, 65 mM sorbitol, 100 mM KCl, 10 mM K_2_HPO_4_, 50 μM EDTA, 1 mM MgCl_2_, 5 mM glutamate, 5 mM malate, 1 mg/ml fatty acid free BSA, and 1 mM ADP)(34). Sample were each divided into six, 1 ml tubes, three for treatment with 2 μM RHPS4 and three without treatment. A dNTP mix of 30 μL each of 10 mM dCTP, 10 mM dGTP, and 10 mM dTTP, as well as 12 μL α-^32^P-dCTP (3000 mmol/Ci), was then added at 17 μL per reaction. Samples were gently rotated at 37°C, with samples collected at 30, 60, 90, and 120 min, spun down, and lysed in proteinase K/SDS buffer and total DNA isolated as described above. DNA was resuspended in 50 μL of TE buffer with RNase A and dimethylurea antioxidant overnight, 25 μL digested with Sac I restriction enzyme, and resolved on 0.6% agarose in 0.5X TBE for 2 hr at 100V/hr. DNA within the gel was nicked, denatured and transferred onto membrane, UV crosslinked, and exposed to phosphorimager for 24 hours. Label incorporation into full-length mtDNA was determined using ImageQuant software package (GE Healthcare). Total mtDNA was determined by random hexamer probing of full-length mtDNA.

### Live cell imaging

To detect the RHPS4 endogenous fluorescence, HeLa cells were seeded on 35mm glass bottom dishes (MatTek Corporation, Ashland, MA) and incubated with 1 μM RHPS4 overnight prior to imaging. The dish was inserted in a closed, thermo-controlled (37°C) stage top incubator (Tokai Hit Co., Shizuoka-ken, Japan) atop the motorized stage of an inverted Nikon TiE fluorescent microscope (Nikon Inc., Melville, NY) equipped with a 60X oil immersion optic (Nikon, CFI PlanFluor, NA 1.43) and NIS Elements Software (Nikon). RHPS4 was excited using either the 470 nm or the 555 nm line of a Lumencor diode-pumped light engine (SpectraX, Lumencor Inc., Beaverton, OR) and detected using ET-GFP and ET-TRITC emission filter (Chroma Technology Corp., Bellows Falls, VT) and ORCA-Flash4.0 sCMOS camera (HAMAMATSU Corporation, Bridgewater, NJ).

For the RHPS4 mitochondrial localization, HeLa cells were treated in culture with 1 μM RHPS4 overnight, washed with PBS, and incubated with MitoTracker Deep Red FM (ThermoFisher Scientific) probe in the dark for 15 min. The RHPS4 signal was detected using 561 nm excitation and TRITC emission filters. Images were captured on a Nikon Ti equipped with a 60X (1.4 NA) optic using a Sweptfield confocal scanhead (35 micron slit; Prairie Instruments, Middleton, WI), a Photometrics Evolve 512 camera (Tucson, AZ) and NIS Elements software. Binary masks were generated in NIS Elements and binary math used to calculate the volume of protein within the mitochondria compartment, as defined by the MitoTracker Deep Red signal. For the 3D rendering of RHPS4 co-localization with mitochondria, live cells were imaged using a 60X (1.40 NA) objective on a Nikon Ti inverted microscope equipped with a Tokai-Hit environmental chamber and Hamamatsu Flash 4.0 CMOS camera and processed in Nikon NIS Elements software.

To assess co-localization of RHPS4 with mitochondria after uncoupling, 143B ρ^+^ and ρ^0^ cells were seeded on 35mm glass bottom dishes and incubated overnight with 1 μM RHPS4 prior imaging. The cells were mounted in a Tokai-Hit environmental chamber atop the stage of a Nikon Ti inverted microscope equipped with a Sweptfield confocal head and imaged using a 60x 1.40 N.A. objective, and Photometrics Evolve EMCCD camera. Cells were imaged at 30 second intervals for fifteen minutes to establish a baseline level of fluorescent signal and observe the amount of photobleaching due to imaging conditions. Immediately after baseline levels were established, 1 μM trifluoromethoxy carbonylcyanide phenlhydrazone (FCCP) compound was added to the culture media and the cells were observed for 15 minutes at 30 second intervals with the same imaging conditions as in the baseline experiment. Data were processed using NIS Elements software.

### Fixed cell microscopy

MEF cells were grown on microscope coverglass (Fisher Scientific) overnight and incubated with 30 μM RHPS4 (Tocris Bioscience) for 30 min prior permeabilization with ice cold methanol for 5 min. Images were taken using a Nikon A1 Spectral Confocal Laser Microscope System and processed in Nikon NIS Elements software (Nikon).

### FRET melting assay

RHPS4 stabilization and selectivity for G4 DNA was determined using FRET assay with F21D as previously described (35). In brief, F21D was annealed 24 h before the FRET study by heating the 0.25 μM F21D to 95 °C for 10 min in 5K buffer (10 mM Li cacodylate pH 7.2, 5 mM KCl, and 95 mM LiCl) then slowly cooled to room temperature over three hours and stored at 4 °C overnight. RHPS4 was added in the concentration range from 0.4 to 4.0 μM bringing the final F21D concentration to 0.2 μM. Samples were thoroughly mixed and equilibrated at least for one hour before FRET measurements. Fluorescence measurements taken every 1 °C from 15 °C to 95 °C, with temperature rate of 1 °C/min on a MJ research Chromo4 (now Bio-Rad, Hercules, CA). Melting temperature was determined from the first derivative of melting data. For FRET-based competition studies, 0.2 μM F21D sample incubated with 1.6 μM RHPS4 was supplemented with 4.8 to 96.0 μM calf thymus (CT) DNA as non-specific, double stranded DNA competitor. Negative control experiment included 0.2 μM F21D with 16 μM CT in the absence of ligand. The errors associated with Tm determination were ± 0.5 °C. All samples were measured in technical duplicate per experiment and each experiment was repeated three times.

### UV-Vis binding studies

UV-Vis studies were performed to demonstrate direct ligand-DNA interaction between mtDNA oligo A or oligo B and RHPS4 using a Cary 300 Varian spectrophotometer with a Peltier-thermostated cuvette holder (error of ± 0.3 °C) in 5K buffer. The advantage of this experiment over FRET assay is the absence of fluorescence tag on DNA. Titrations were performed by stepwise additions of annealed oligo A or oligo B to a 0.3 mL solution of 20 μM RHPS4 in a 1 cm quartz cuvette. Resulting solution was mixed and equilibrated for two min. UV-Vis spectra were acquired in 350 – 670 nm range. Titration was deemed complete when the spectra collected after three successive additions of G4 oligonucleotides were nearly superimposable. At this point, typical [G4]/[RHPS4] was ~ 3. Data were treated as described in detail in our earlier work and spectra were corrected for dilution effect (36). Direct fitting of UV-Vis data (assuming two-state equilibrium) was used to obtain the values of binding constant, Ka. Hypochromicity (% H, decrease in signal intensity) and red shift (change in peak position) were extracted from UV-Vis data as described in (36). Titrations were repeated at least three times.

#### Thermal Difference Spectra (TDS)

<u>UV-VIS TDS</u> were obtained by taking the difference between UV-VIS signal of DNA above and below its melting point, customarily at 4 and 90 °C in three buffers: 10Li (10 mM lithium cacodylate buffer, pH 7.2), 5K (10Li supplemented with 5 mM KCl and 95 mM LiCl), and 100Li (10Li supplemented with 100 mM LiCl). Characteristic signature of G4 structures in TDS consists of a negative peak at ~295 nm and two positive peaks at ~243 and ~273 nm (37).

### CD experiments

CD wavelength scans were collected on each sample to determine the extent and type of the folding in 10Li, 5K, or 100Li. 5K buffer was used unless otherwise indicated. The experiments were performed using AVIV 410 or 435 spectrometer equipped with a Peltier heating unit, with the following parameters: 2 nm bandwidth, 220-330 nm window, 1 sec average time, 10 °C temperature, and 3 scans. Resulting data were treated as described in our previous work (36). CD melting experiments were performed by heating the samples from 4 to 90 °C and then back to test reversibility of the melting process. CD melting was performed using the following parameters: wavelength, 264 nm; bandwidth, 4 nm; T range: 4 - 95 °C; T step: 1 °C; dead band: 0.33 °C; equilibrating time, 0.5 min; averaging time, 20 - 30 sec. The B+RHPS4 was monitored in 100Li and 5K buffers at 295 nm instead of 268 nm because of its intense antiparallel character. CD scans were collected after completion of the CD melt to assure that the samples have returned to equilibrium. Melting transitions for oligo A and B, and their complexes with RHPS4, were accompanied by large hysteresis and thus were not reversible. Melting temperature, designated T_1/2_, obtained from such data is only relevant for the specified conditions. On the other hand, melting of m.10191T, m.10191T>C was nearly reversible (hysteresis 2-6 °C); thus the data were fit using a two-state model with constant enthalpy, ΔH = const (zero heat capacity, Cp = 0) (38). Melting experiments were repeated at least three times.

### PCR stop assay

Polymerase stop assay was run as reported (27). Briefly, the experiment was conducted adding RHPS4 (0; 1; 2; 5; 10 μM) to 25 μl reaction containing DreamTaq DNA Polymerase (ThermoFisher Scientific), 4.6 ng/reaction DNA, 0.2 mM dNTPs and 0.4 μM of each primer. The PCR was performed in a ProFlex thermal cycler (ThermoFisher Scientific) with the following PCR amplification profile: one cycle of 95°C for 2 min; 30 cycles of 95°C for 30 sec, 56°C for 30 sec, and 72°C for 30 sec. Products of amplification were run on a 2% agarose gel and detected with ethidium bromide fluorescence on a Bio-Rad ChemiDoc MP Imaging System (Bio-Rad, Carlsbad, CA). Quantitation was performed on three or more independent experiments.

### Western blotting analysis

Cell pellets were lysed in RIPA buffer (50 mM Tris-HCl, pH 7.4, 150 mM NaCl, 0.25% sodium deoxycholate, 1 mM EDTA, 1% NP-40, 1X Complete protease inhibitor cocktail [Roche Molecular Diagnostics, Pleasanton, CA], and phosphatase inhibitor cocktail [ThermoFisher Scientific]). Cell extracts were separated on a 4-12% Bis-Tris polyacrylamide gel and proteins analyzed by immunoblotting using primary antibodies against the following proteins: ATP5A, UQCRC2, MTCO1, SHDB and NDUFB8 (Total OXPHOS Rodent WB Antibody Cocktail; Abcam, Cambridge, MA); SDHA (Abcam); TFAM (PhosphoSolutions, Aurora, CO); and GAPDH (EMD-Millipore, Billerica, MA) followed by IR Dye-labeled secondary antibodies (Li-Cor, Lincoln, NE). Signal intensity was detected with the Li-Cor Odyssey infrared imager at 680 and 800 nm and processed with manufacturers software.

### Transcript analysis

To quantify gene expression by real-time quantitative polymerase chain reaction (qRT-PCR), RNA was isolated from cell pellets with RNeasy Mini Kit (Qiagen, Germantown, MD) and genomic DNA contamination was removed using DNA-free DNA Removal kit (ThermoFisher Scientific). Quality of the extracted RNA was assessed by 1% agarose gel electrophoresis and from the A_260nm_/A_280nm_ absorbance ratio (Nanodrop 1000, ThermoFisher Scientific). Next, cDNA was synthesized using the High Capacity cDNA Kit (ThermoFisher Scientific) with either random (provided by manufacturer) or specific primers (see Primers and probes below). Finally, processed and unprocessed RNA expression levels were assessed using the ΔΔC_q_ method (31). The primer/probe sets were designed and their compatibility validated *in silico*. A serial dilution approach was used to validate the multiplex reaction. All qPCR reactions were carried out on either a StepOnePlus or QuantStudio 5 thermal cycler (ThermoFisher Scientific). All experiments were run in triplicate and the gene expression levels normalized to the B2M results.

### RNA-Seq analysis

For RNA-Seq experiments, total RNA concentration and quality from each sample was assessed using Qubit 2.0 fluorometer (ThermoFisher Scientific) and Agilent TapeStation 2200 (Agilent, Santa Clara, CA). Total RNA libraries were generated using Illumina TruSeq Stranded Total RNA Sample Preparation Guide Rev. E (Illumina, San Diego, CA). In brief, this process depletes nuclear ribosomal RNA (rRNA) and then fragments the remaining RNA, which was converted into first strand cDNA using reverse transcriptase and random primers. The second strand cDNA was generated using DNA polymerase I and RNase H. The cDNA fragments were further processed for adapter ligation, and then enriched with PCR to create the final cDNA library.

The cDNA libraries were validated using Illumina-compatible DNA primers (KAPA Biosystems, Wilmington, MA) and Qubit 2.0 fluorometer. Quality was examined using Agilent Tapestation 2200. The cDNA libraries were pooled at a final concentration 1.8 pM. Cluster generation and 75 bp paired read dual-indexed sequencing was performed on Illumina NextSeq 500.

Quality control for raw fastq files were performed with FastQC (39), the low quality reads and 3’ adapters were trimmed with Trim Galore! and Cutadapt (40, 41). The trimmed reads were aligned to reference genome (mm10) with the RNA-seq aligner STAR (42), and the resultant bam files were converted to bigWig format using the *bam2wig* function in RSeQC (43) for the visualization of read coverage in genome browser, such as IGV (44). Subsequently, gene expressions in each sample were quantified as the number of read fragments that were uniquely mapped to genes using featureCounts (45). The raw count was then normalized as FPM (Fragments Per Million mapped) to eliminate the influence of sequencing depth. Gene differential expression analysis was performed on the raw count table using DESeq2 (46). Genes with FDR < 0.01 and fold-change > 1.5 were determined as differentially expressed genes (DEGs). Enrichment of the DEGs in KEGG pathways was performed with Fisher’s exact test.

### Measurement of Oxygen Consumption Rate (OCR)

OCR of control and patient primary human fibroblasts treated with 0 or 1 μM RHPS4 for 48 h was measured using the XF-24 Seahorse system (Seahorse Bioscience, Billerica, MA). After 48 h incubation with vehicle (DMSO) or 1 μM RHPS4, cells were washed with pre-warmed assay medium and seeded in XF 24-well cell culture microplates (Seahorse Bioscience) at 5 × 10^3^ cells/well, five wells per group. The basal respiration data were collected and the cells sequentially treated with 1 μM oligomycin, 150 μM dinitrophenol, and 160 nM rotenone as previously optimized for this system (47). All experiments were performed in unbuffered DMEM supplemented with 5 mM glucose. Cell lines used were: normal human primary fibroblast; patient fibroblast carrying ~50% heteroplasmic mutation load for m.13513G>A; patient fibroblast carrying ~90% mutation load for m.10191G>C.

### Primers and probes

The following primers were used for in vitro G-quadruplex formation assays:

F21D (5’-6-FAM-G_3_(TTAG_3_)_3_-Dabcyl-3’);
Oligo A (5’-GGATGGGGTGGGGAGG-3’);
Oligo B (5’-GGGGGATGCGGGGG-3’)

Primers used to perform the PCR stop assay were:

mt ND3 primer 1 (5’-AAAATCCACCCCTTACGAGT-3’)
mt ND3 primer 2 (5’-TATTGGCTAAGAGGGAGTGG-3’)
mt COX1 primer 1 (5’-GGTTCGATTCCTTCCTTTTT-3’)
mt COX1 primer 2 (5’-GCCTGACTGGCATTGTATTA-3’)
mt non G4 primer 1 (5’-GCACTCGTAAGGGGTGGAT-3’)
mt non G4 primer 2 (5’-TCGAGTCTCCCTTCACCATT-3’)

The specific primers used for the cDNA synthesis were:

E-ND6/ND6 (5’-TCCAAACACAACCAACATCC-3’);
F-RNR1-V-RNR2 (5’-GGTGTAGGCCAGATGCTTTAAT-3’);
ND1-IQM (5’-AAGAGGGCTTGAACCTCTATAA-3’);
ND6-E-Cytb (5’-GGCAGGTAGGTCAATGAATGAGTG-3’);
B2M (5’-CCGTTCTTCAGCATTTGGATTT-3’).

For the mature RNA assay, the mouse probes ND6, RNR1, RNR2, ND1, ND2, COX1, COX2, ATP6, COX3, ND3, ND4L, ND4, ND5, CYTB were from ThermoFisher Scientific. The primers and probes used for the parental RNA assays were purchased from Integrated DNA Technologies (IDT):

E-ND6 probe (5’-/56-FAM/TTGGTTGGT/ZEN/TGTCTTGGGTTAGCA-3’)
E-ND6 primer 1 (5’-GTCATTGGTCGCAGTTGAATG-3’)
E-ND6 primer (5’-ACCTCCATAAATAGGTGAAGGC-3’)
F-RNR1-V-RNR2 probe (5’-/56-
FAM/AAACACAAA/ZEN/GGTTTGGTCCTGGCC/-3’)
F-RNR1-V-RNR2 primer 1 (5’-GCTTAATAACAAAGCAAAGCACTG-3’)
F-RNR1-V-RNR primer 2 (5’-TCTATGGAGGTTTGCATGTGTA-3’)
L-ND1 probe (5’-/56-FAM/AGGATTTGA/ZEN/ACCTCTGGGAACAAGGT/-3’)
L-ND1 primer 1 (5’-AGCCAGGAAATTGCGTAAGA-3’)
L-ND1 primer 2 (5’-GGGACGAGGAGTGTTAGGATA-3’)
ND6-E-CYTB probe (5’-/56-FAM/TTGGTTGGT/ZEN/TGTCTTGGGTTAGCA/3’)
ND6-E-CYTB primer 1 (5’-AAACAACCAACAAACCCACTAAC-3’)
ND6-E-CYTB primer 2 (5’-GCAGTTGAATGCTGTGTAGAAATA-3’)
B2M probe (5’-/5HEX/TTCAAGTAT/ZEN/ACTCACGCCACCCACC-3’)
B2M primer 1 (5’-ACGTAGCAGTTCAGTATGTTCG-3’)
B2M primer 2 (5’-GGTCTTTCTGGTGCTTGTCT-3’)

For the mature RNA assay, standard qPCR thermal parameters were used: one cycle of 95 °C for 20 sec then 40 cycles of 95 °C for 1 sec and 60 °C for 20 sec. For the parental RNA assays, the optimized conditions were as follows: L-ND1 or F-RNR1-V-RNR2 mouse genes expression, one cycle of 95°C for 10 min then 40 cycles of 95°C for 20 sec, 56°C for 20 sec and 60°C for 20 sec; ND6-E-CYTB or E-ND6 mouse gene expression, one cycle of 95°C for 10 min then 40 cycles of 95°C for 20 sec, 60°C for 20 sec and 60°C for 20 sec.

### Statistical analysis

Statistical analysis was performed by one-way ANOVA with Dunnett’s posthoc analysis or Student two-tailed test using GraphPad Prism software (GraphPad, La Jolla, CA). P-values of <0.05 were accepted as a significant difference. All data are expressed as mean +/-standard error of the mean (SEM).

## RESULTS

### Identification of RHPS4 as a mitochondrial G4 ligand that induces hallmarks of mtDNA replication pausing

Our objective was to stabilize G4 structures in mitochondria to determine their impact on mitochondrial DNA, gene expression, and respiratory function. In the nucleus, ligand-mediated increases in G4 stability are known to cause replication pausing (9). By extension, we expected that small molecules able to increase G4 stability in mitochondria would also cause replication pausing, leading to mtDNA depletion (26). We tested four known G4 ligands (Braco-19, NMM, Phen-DC3, and RHPS4) for their ability to induce mtDNA depletion in cultured mouse embryonic fibroblast cells (Figure 1A). Because mitochondrial localization of small molecules has been observed with both neutral and positively charged lipophilic compounds, these compounds were selected as representatives of varied ranges of scaffold structure and charge. The positively charged compound RHPS4 induced significant and reproducible depletion of mtDNA content with mild effect on cell growth (Figure 1B). We further established that the decrease of mtDNA levels was time-dependent (Figure 1C) and reversible (Figure 1D). Supporting the notion that stabilization of G4 structures would impede replication, we found that RHPS4 treatment reduced the ratio of linear to circular mtDNA content (Figure 1E and F), a feature previously observed in ddC-induced mtDNA replication pausing and depletion (32).

**Figure 1.**
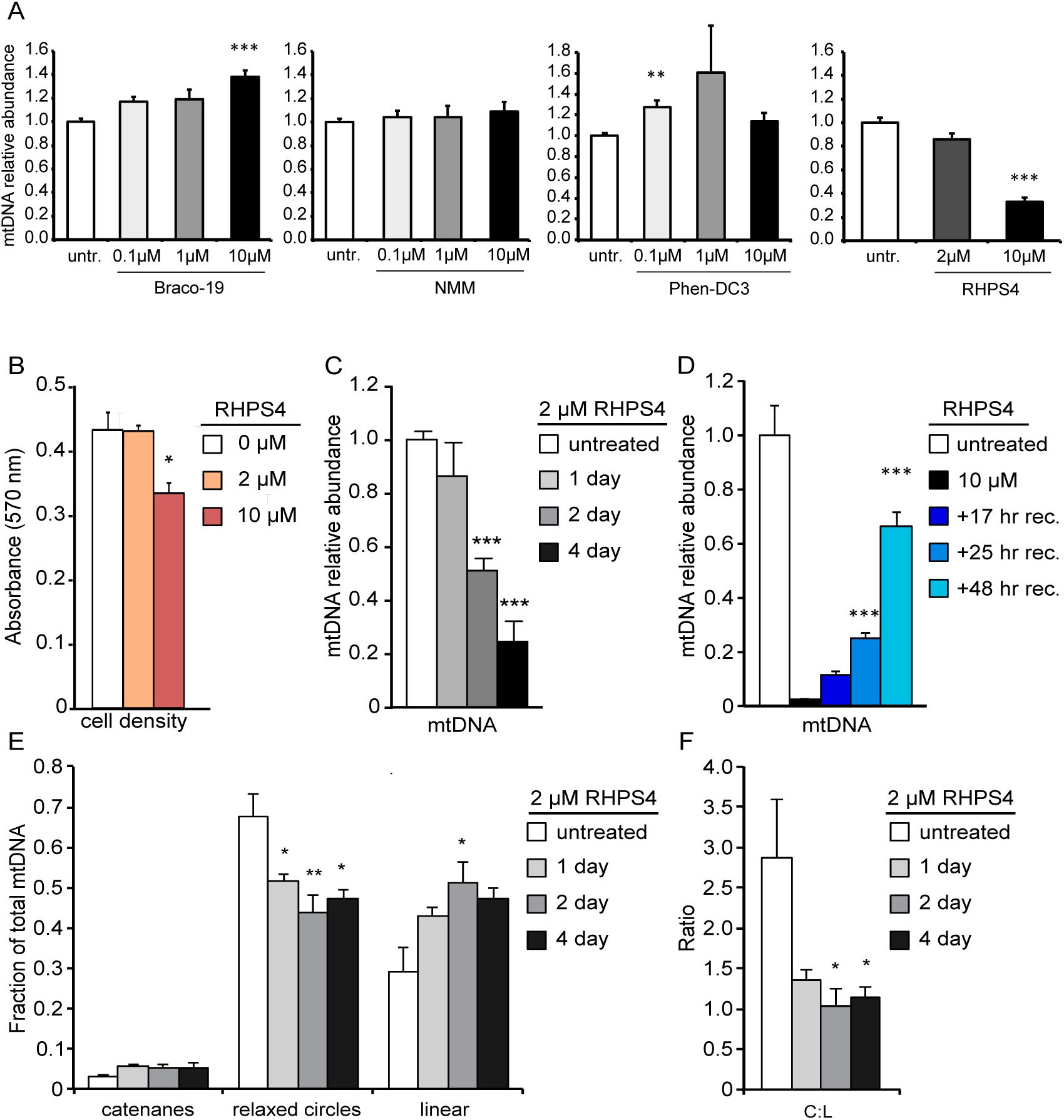
G4-ligand exposure causes reversible mtDNA depletion. (A) Determination of MEFs sensitivity to different G-quadruplexes ligands. mtDNA depletion analyzed by qPCR in WT MEFs cultured in presence of Braco-19, NMM, Phen-DC3, or RHPS4 at the indicated concentrations for 24 hours (untr.; untreated). Each treatment set is normalized to untreated control. All data are mean +/- SEM (n=4; p-values calculated by one-way ANOVA with Dunnett’s posthoc analysis: **<0.01, ***<0.001). (B) Crystal violet assay on mouse embryonic fibroblasts (MEFs) treated with RHPS4. (C) Time-dependent mtDNA depletion as analyzed by qPCR. (D) Reversibility of mtDNA depletion in WT MEFs. Cells were exposed to 10 μM RHPS4 for 4 days and followed by recovery (rec.) for the indicated time. (E) Alterations in the relative abundance of mtDNA catenanes, relaxed circles, and linear molecules from panel B time course. (F) RHPS4 decreases the mtDNA circular: linear ratio (C:L), a replication pausing hallmark. All data are mean +/- SEM (n=3-8; p-values calculated by one-way ANOVA with Dunnett’s posthoc analysis: *<0.05, **<0.01, ***<0.001).

We tested whether RHPS4 was acting directly on mtDNA to block replication using an *in organello* replication assay (34, 48) (Figure 2). Using isolated mouse liver mitochondria, 2 μM RHPS4 inhibited both radiolabel incorporation into full-length mtDNA (Figure 2A) and total full-length mtDNA synthesis (Figure 2B). Consistent with the idea that RHPS4 binds to mtDNA G4 sequences to block DNA synthesis, we found that 10 μM RHPS4 interferes preferentially with the synthesis of mtDNA G4 sequences (ND3 and COX1) relative to non-G4 forming sequences in a PCR stop assay of total DNA isolated from HeLa cells (Figure 2D,E). These data strongly suggest that RHPS4 causes mtDNA replication defects through direct interaction with mitochondrial nucleic acids.

**Figure 2.**
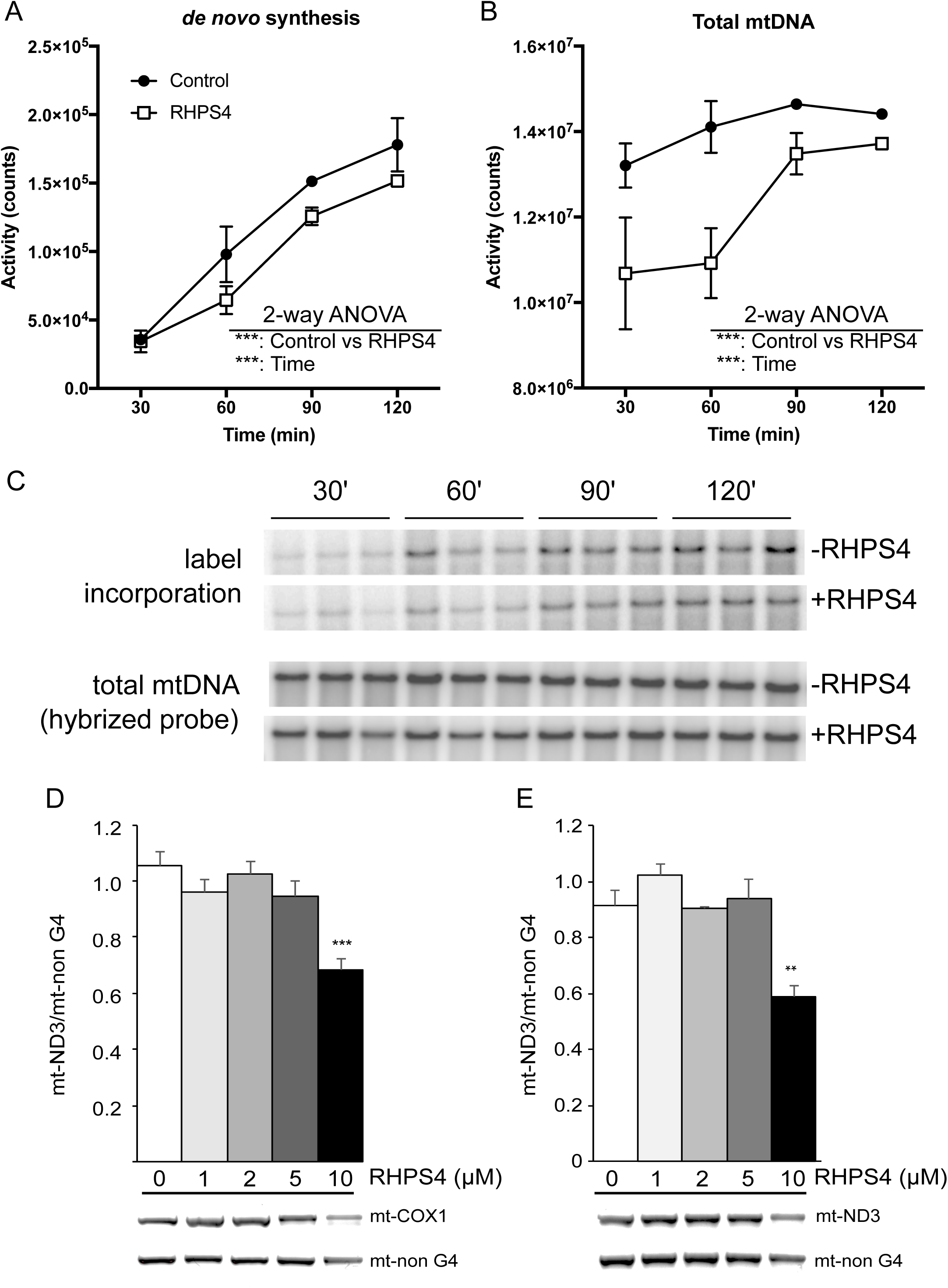
RHPS4 inhibits mtDNA replication or polymerization *in vitro*. (A,B) DNA replication in isolated mouse liver mitochondria with or without 2 μM RHPS4. (A) Radiolabel incorporation into full-length mtDNA across 30-120 min timecourse. (B) Total full-length determined by hybridization probing and autoradiograpy. (C) Raw radiographic images of full length mtDNA. (D,E) PCR stop assay using untreated HeLa cellular DNA amplifying mt-COXI (G4 amplicon; D) or mt-ND3 (G4 amplicon; E) and mt-non G4 (no G4 in amplicon). Amount of RHPS4 in PCR reaction as indicated. Data are shown as mean +/- SEM (n=3). P-values determined by 2-way (A,B) or 1-way ANOVA (D,E; *<0.05; **<0.01; ***<0.001).

We next investigated the subcellular localization of RHPS4 using live cell imaging (Figure 3). Previous reports indicated that innate fluorescence of RHPS4 localizes to the nucleus after incubation with 30 μM compound for 30 min followed by alcohol permeabilization (49) or with 20 μM for 24 hours in live cells (50). RHPS4 has spectral characteristics compatible with TRITC and FITC channels in live cell imaging, which allows equilibration of the molecule and imaging without fixation. In our hands, 1 μM RHPS4 overnight exposure was sufficient to achieve fluorescence in these channels without signal in the far-red wavelengths (Figure 3A). To determine whether the TRITC population localized to mitochondria, we co-stained mitochondria with MitoTracker Deep Red and acquired live cells images in the TRITC and far red channels (Figure 3B). The RHPS4 fluorescence pattern was not altered by the addition of MitoTracker, and 100% of the RHPS4 volume was positive for MitoTracker Deep Red. Similarly, 96% of the mitochondrial volume (defined by MitoTracker) contained RHPS4. Strongly enriched mitochondrial localization can be observed in three-dimensional renderings of RHPS4 and MitoTracker Deep Red with no discernible nuclear staining in live cells using LiveScan Swept Field confocal microscopy (Figures S1). Unfortunately, attempts to co-localize RHPS4 with mitochondrial sub-compartments using *in situ* immuno co-localization of fixed samples were unsuccessful due to RHPS4 diffusion during processing.

**Figure 3.**
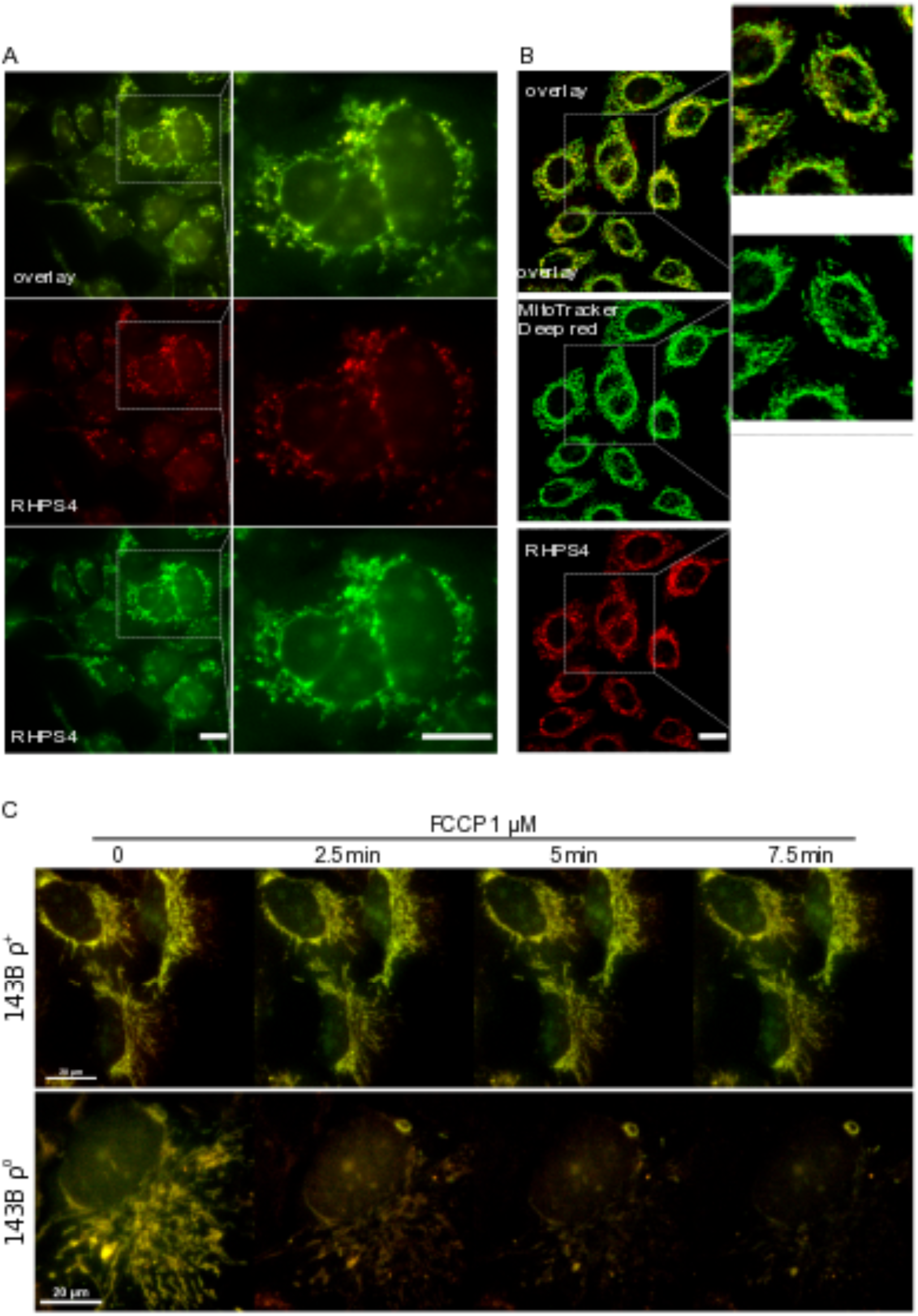
RHPS4 localizes to mitochondria by membrane potential and is retained by binding nucleic acids. (A) Live cell widefield images of HeLa cells incubated with 1 μM RHPS4 overnight. The images show RHPS4 endogenous fluorescence from FITC channel (green), TRITC channel (red), and channel overlay (yellow). Some RHPS4 localizes to the nucleolus with extended light source exposure. (B) Single confocal plane images of live cells treated with RHPS4 detected in the TRITC channel (red), co-stained by MitoTracker Deep Red (green) and their merge (yellow). 100% of RHPS4 value was MitoTracker Deep Red positive, whereas MitoTracker Deep Red was 96% RHPS4 positive. No fluorescence was detected when both compounds were absent. Digital zoom of images shown on right of each panel. Scale bars in panels A and B are 10 μm. (C) Time course of live cell Sweptfield confocal images of 143B ρ^+^ (upper) and 143B ρ^0^ (lower) cells incubated with 1 μM RHPS4 overnight. After the medium was changed, the cells were treated with 1 μM FCCP and multiple individual fields imaged over 15 min. Overlay (yellow) of the endogenous RHPS4 fluorescence detected in the FITC (green) and TRITC (red) channels. Scale bars in panel C are 20 μm.

Previous localization to the nucleus is potentially non-physiological. We find a limited localization to nucleoli was apparent only after extended exposure to epifluorescence light sources (demonstrated in Figure S2), a localization pattern previously reported for RHPS4 in live cell localization studies using wide field fluorescence (50). Rapid image acquisition with Swept field confocal prevented nucleolar localization as light exposure induced a change in localization (as in Figure 3B and Figure S1). We also found that a combination of nucleolar and diffuse cellular localization occurred in MEF cells after exposure with 30 μM RHPS4 for 30 min and methanol fixation (Figure S3A), a method also previously reported (49). Taken together, we provide strong evidence for a mitochondrial localization of RHPS4 at low doses in live, non-cancerous cells, and we can recapitulate non-mitochondrial RHPS4 localization by both photoactivation or fixation artifacts.

To investigate the mechanism of RHPS4 localization, we tested whether mitochondrial membrane potential or the permeability transition pore (PTP) were essential for RHPS4 uptake. Numerous positively charged compounds localize to mitochondria via membrane potential, which has been used to target small molecules to the mitochondrial compartment through moieties such as triphenylphosphonium (TPP)(51). The targeting of TPP-coupled molecules to mitochondria is blocked by dissipation of the mitochondrial membrane potential. We first preloaded 143B ρ^+^ (normal mtDNA) or ρ^0^ (devoid of mtDNA) cells with 1 μM RHPS4 overnight prior to dissipating the mitochondrial electrochemical gradient with the mitochondria-specific uncoupler FCCP. After confirming the photostability of the RHPS4 under short repeated measurements (Figure S3B), 1 μM FCCP final concentration was applied to live cells and multiple fields were imaged in 2.5 min time increments (Figure 3C). We observed a loss of RHPS4 mitochondrial localization after FCCP treatment only in 143B ρ^0^ cells. Unfortunately, an experiment using the reverse exposure order (FCCP then RHPS4) cannot be performed because the cells cannot be maintained in FCCP overnight. As well, the potential G4 ligand BMVC was suggested to localize to mitochondria through the mitochondrial PTP (27), which is blocked by 1 μM cyclosporin A (CsA). We found that RHPS4 localization was not impacted by 1 μM CsA pretreatment (Figure S4).

### Selective binding of RHPS4 to G4 structures

We next confirmed RHPS4 preferential binding of G4 structures vs. double-stranded (ds) DNA, which is crucial to the interpretation of the biological impact of G4 structures stabilization. Toward that end, a well-defined, G4-forming fluorescently labeled human telomere sequence, F21D, was subjected to fluorescence resonance energy transfer (FRET)-melting assays (35) with RHPS4. We found that sub-micromolar RHPS4 concentrations were sufficient to increase the melting temperature of F21D by at least 15 °C (Figure 4A). This stabilization was poorly competed by 480 equivalents of non-specific dsDNA (Figure 4B). These data demonstrate the strong selectivity of RHPS4 for G4 structures over dsDNA and also establish that RHPS4 binding thermodynamically stabilizes G4 structures, consistent with previous reports (52–54).

**Figure 4.**
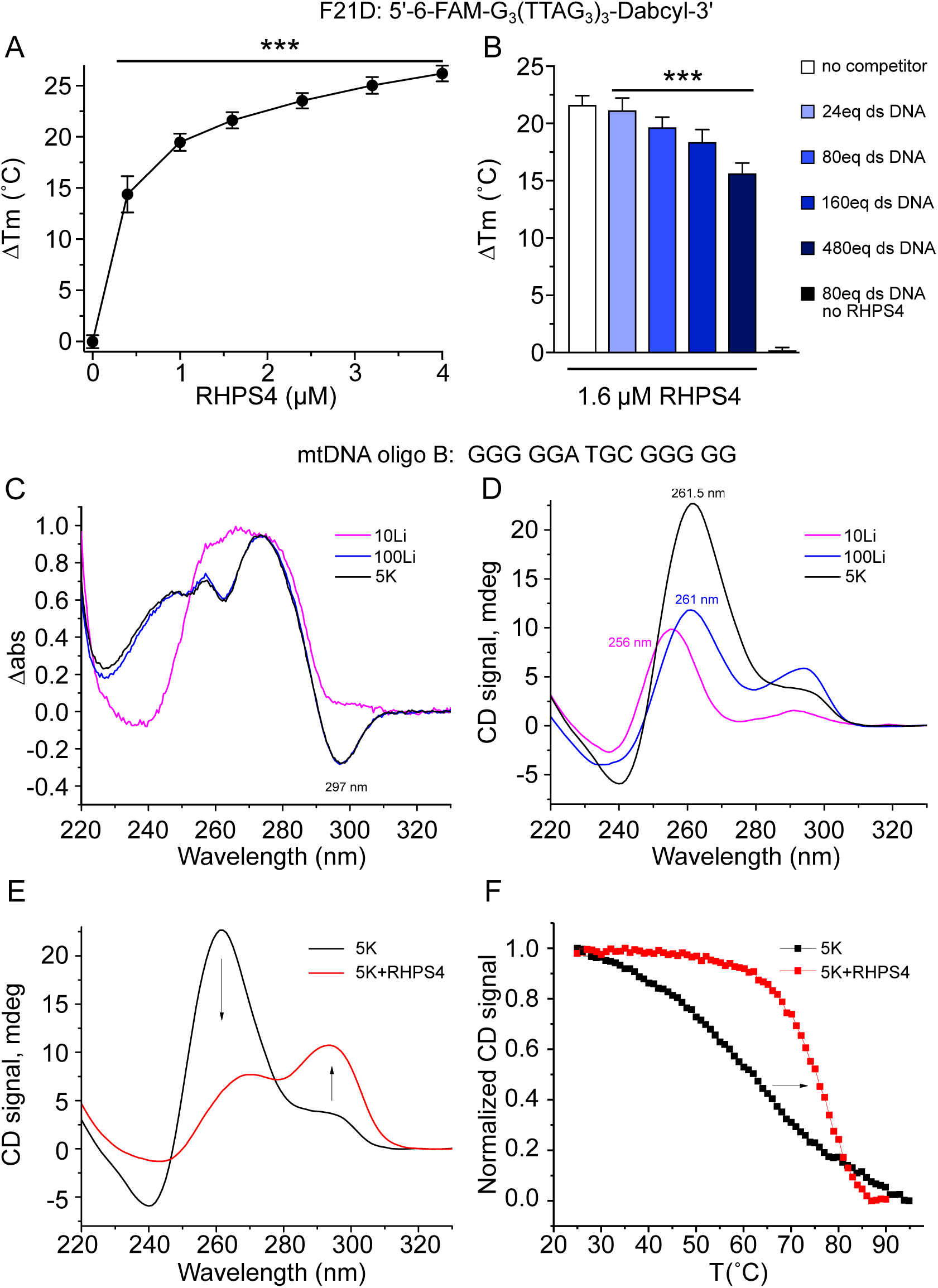
RHPS4 is a G-quadruplex ligand that stabilizes antiparallel structures in G4-forming mtDNA sequences. (A) FRET-melting profile of 0.2 μM F21D, a fluorescently labeled G4-forming human telomeric DNA, in the presence of increasing concentration of RHPS4. (B) FRET-melting competition assay showing the inability of dsDNA to compete with F21D for RHPS4 binding. Negative control consists of 0.2 μM F21D, 80 eq of dsDNA and no ligand. P-values for Panel A and B were calculated by one-way ANOVA with Dunnett’s posthoc analysis: ***<0.001. (C) UV-VIS TDS spectra of oligo B; characteristic negative peak at 295 nm supports G4 structure formation in 100Li or 5K buffer; this peak is absent in 10K buffer. (D) CD spectra of 5 μM oligo B in 10Li, 100Li, or 5K buffers. Parallel and antiparallel G4 character is detected at ~264 nm and ~295 nm, respectively. (E) CD spectra of 5 μM oligo B in the presence (red lines) or absence (black lines) of 10 μM RHPS4 in 5K buffer. (F) CD melting curve for oligo B, with or without RHPS4. Signal was monitored at 260 nm for oligo B or 295 nm for oligo B in the presence of RHPS4.

### Interactions between RHPS4 and G4 structures in mtDNA sequences

We previously demonstrated that mtDNA oligo A and B sequences form predominantly parallel G4 conformations as signified by the peak at 262 nm in circular dichroism (CD) spectra in the presence of 5 mM K^+^ (6). As part of characterizing interactions of RHPS4 with mtDNA sequences, we tested the ion dependence of G4 structure formation of the se sequences. TDS and CD signal of oligo A or oligo B in 5K or 100Li buffer, but not 10Li buffer, are consistent with G4 structure (Figure 4C,D and Figure S5A,B). The addition of 5 μM RHPS4 in 5K buffer induced a marked antiparallel conformation in both oligo sequences (Figure S5C and Figure 4E) signified by the increase in peak intensity at 295 nm (and decrease of 264 nm peak); notably, oligo B was more sensitive to the presence of RHPS4. Similar trends were observed without K^+^ in 100Li buffer (Figure S6A,B). Consistent with our data, RHPS4 was previously shown to induce antiparallel G4 conformation in human telomeric DNA in the presence or absence of K^+^ (52).

We next assessed the thermodynamic stability of oligo A and B in the presence of 2 eq of RHPS4 by CD in 5K buffer. RHPS4 increased melting temperatures of G4 structures formed by oligo A and B by 7.4 and 14.4 °C, respectively (Figure 4F and Figure S5D). This stabilization is even more pronounced in 100Li buffer, where RHPS4 at 2 eq stabilizes oligo A and oligo B by 24.2 and 38.4 °C, respectively. Our data are in agreement with the previous observation that RHPS4 stabilized human telomeric DNA and *c-kit* oncogene promoter G4 sequences by ~20 °C (54). In addition to stabilization, the presence of RHPS4 led to a significant decrease in hysteresis, or retardation between heating and cooling kinetics, for oligo A from 28 to 19 °C, (Figure S6C,E). The significant hysteresis in our data suggests that melting is not a simple two-state thermodynamic process, rather it is either kinetically controlled or the system contains multiple species in equilibrium (Figures S6C-E). Decreased hysteresis in the presence of RHPS4 could be due to either faster kinetics of GQ folding in the presence of RHPS4 or selective binding of RHPS4 to one of the multiple conformations that exist at elevated temperatures. Combined, our CD data strongly indicate that RHPS4 preferentially binds, stabilizes, and increases the antiparallel component of sequences with G4-forming potential in the mitochondrial genome both in the presence and absence of K+ ions as also observed for RHPS4 binding to human telomeric DNA (54).

To determine the strength of binding of RHPS4 to oligo A and to oligo B, we performed UV-Vis spectral titrations. Curve fitting yielded binding constants (K_a_) of 0.5 × 10^6^/M, similar for both oligos with [G4]/[RHPS4] ratios of 1:3 for oligo A and 1:2 for oligo B (Figure S7). We also observed significant red shift of 11.5 nm at 510 nm wavelength and enhancement of hyperchromicity by ~22 %, which demonstrate close, direct contact between G4 and RHPS4 (Figure S7E), consistent with end-stacking of RHPS4 onto the terminal G4-tetrad described previously (54).

### RHPS4-mediated respiratory complex depletion occurs through a mitochondrial transcription inhibition mechanism

To determine the impact of RHPS4 exposure on mitochondrial respiratory function, we first examined the abundance of representative subunits of the mitochondrial OXPHOS complexes in MEFs exposed to varying concentrations of RHPS4 (Figure 5). MEFs showed significant subunit depletion from Complexes III, IV, and V at one day of 10 μM RHPS4 exposure, and Complexes III and IV at 2 μM RHPS4. In the absence of mtDNA depletion (2 μM RHPS4), the abundance of Complex II subunits and the mitochondrial DNA binding protein TFAM were not affected. Complex I, which has seven mitochondrially-encoded subunits, was not detected by western blot in MEF cells. To confirm this observation, we repeated the experiment using murine C2C12 myotubes (Figure S8) and found that Complex III, IV and I subunits abundance was decreased, but not TFAM or Complex II subunits (Figure S8A), unless mtDNA levels were depleted (Figure S8B). As with MEF exposure (Figure 1E,F), an increase in the ratio of linear to circular mtDNA forms was observed in myotubes (Figure S8C). These data suggest that RHPS4 effects on mitochondria are general across cell types, not specific to cancer cells, and support the hypothesis that RHPS4 induces gene expression defects at concentrations lower than those required to cause mtDNA depletion.

**Figure 5.**
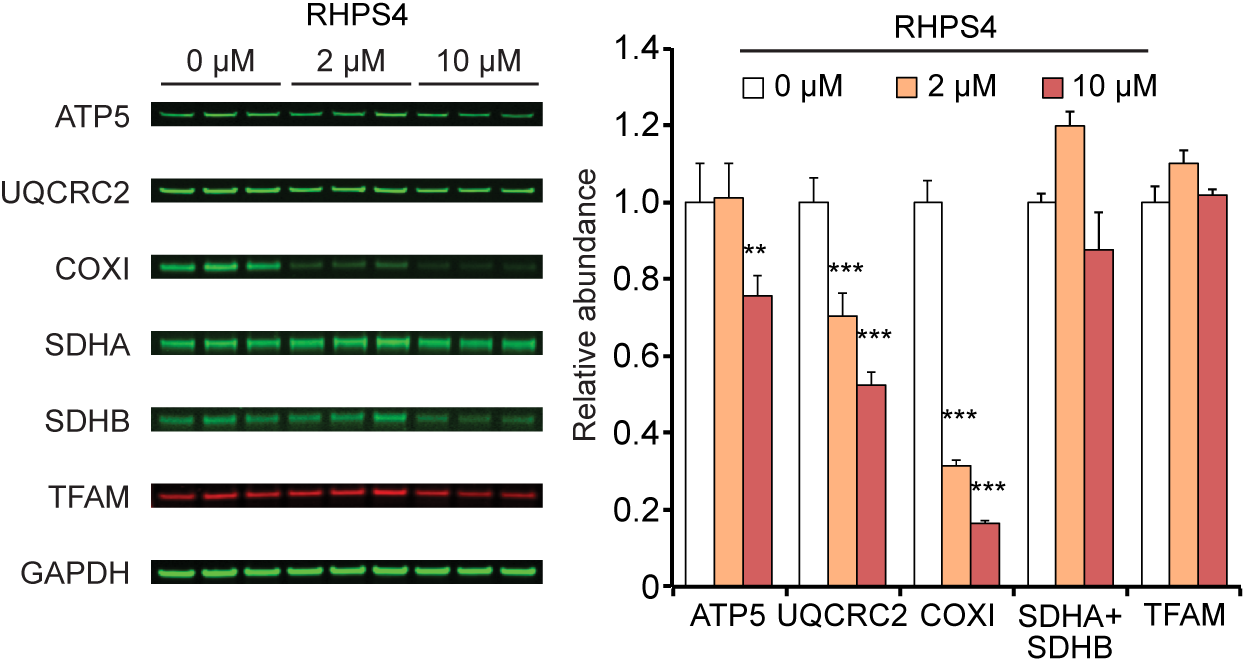
Low-dose RHPS4 exposure causes depletion of representative subunits of complexes III and IV, but not II and V. (LeftFluorescent western blot analysis of OXPHOS proteins in MEFs cultured in presence 0, 2, and 10 μM RHPS4 for 24 hrs. Transferred membranes were probed for ATP5 (Complex V), UQCRC2 (Complex III; CoIII), COXI (Complex IV; CoIV), SDHA and SHDB (Complex II; CoII), TFAM, and GAPDH (loading control). Complex I subunits were not detected in MEFs. (Right) Bar graph of OXPHOS proteins and TFAM normalized to GAPDH and control cell exposure levels (0 μM RHPS4). Shown are mean values +/- SEM (n=3; p-values calculated by one-way ANOVA with Dunnett’s posthoc analysis: **<0.01, ***<0.001).

We next examined alterations in the steady-state transcript levels of mitochondrially-encoded genes. Mitochondrial transcription is initiated by three promoters, the Light Strand Promoter (LSP), Heavy Strand Promoter 1 (HSP1), and Heavy Strand Promoter 2 (HSP2; reviewed in (55)). Transcription initiated at these promoters forms polycistronic RNAs that are then efficiently cleaved and processed into mature RNAs (Figure 6A)(56, 57). Our initial studies used commercial TAQMAN probes to detect the relative abundance of the two ribosomal and the 13 protein coding genes (non-tRNA RNAs) (Figure 6B). We noted that LSP and HSP1-derived transcripts (ND6 and RNR1, respectively) were significantly reduced at RHPS4 exposures that do not cause mtDNA depletion (2 μM); however, we found that RNAs specific to the HSP2 promoter activity showed greater depletion as distance from HSP2 increased (from ND1 to CYTB). The relative abundance of ND6, RNR1, ND1, and CYTB was confirmed by evaluating strand-specific levels of steady-state mitochondrial mRNA in MEFs by RNA-SEQ (Figure S9A). Effects on the HSP2-derived tRNAs loosely recapitulated the promoter distance effect in mRNA (Figure S9B); however, the number of sequencing reads was too low for a robust result across all sequences. We also confirmed the RHPS4-mediated effect on ND6, RNR1, ND1 and CYTB and by qRT-PCR in C2C12 myotubes (Figure S9D). Using human and mouse cultured cells, these data indicate that the stabilization of G4 structures alters mitochondrial RNA abundance without being a secondary effect of mtDNA depletion.

**Figure 6.**
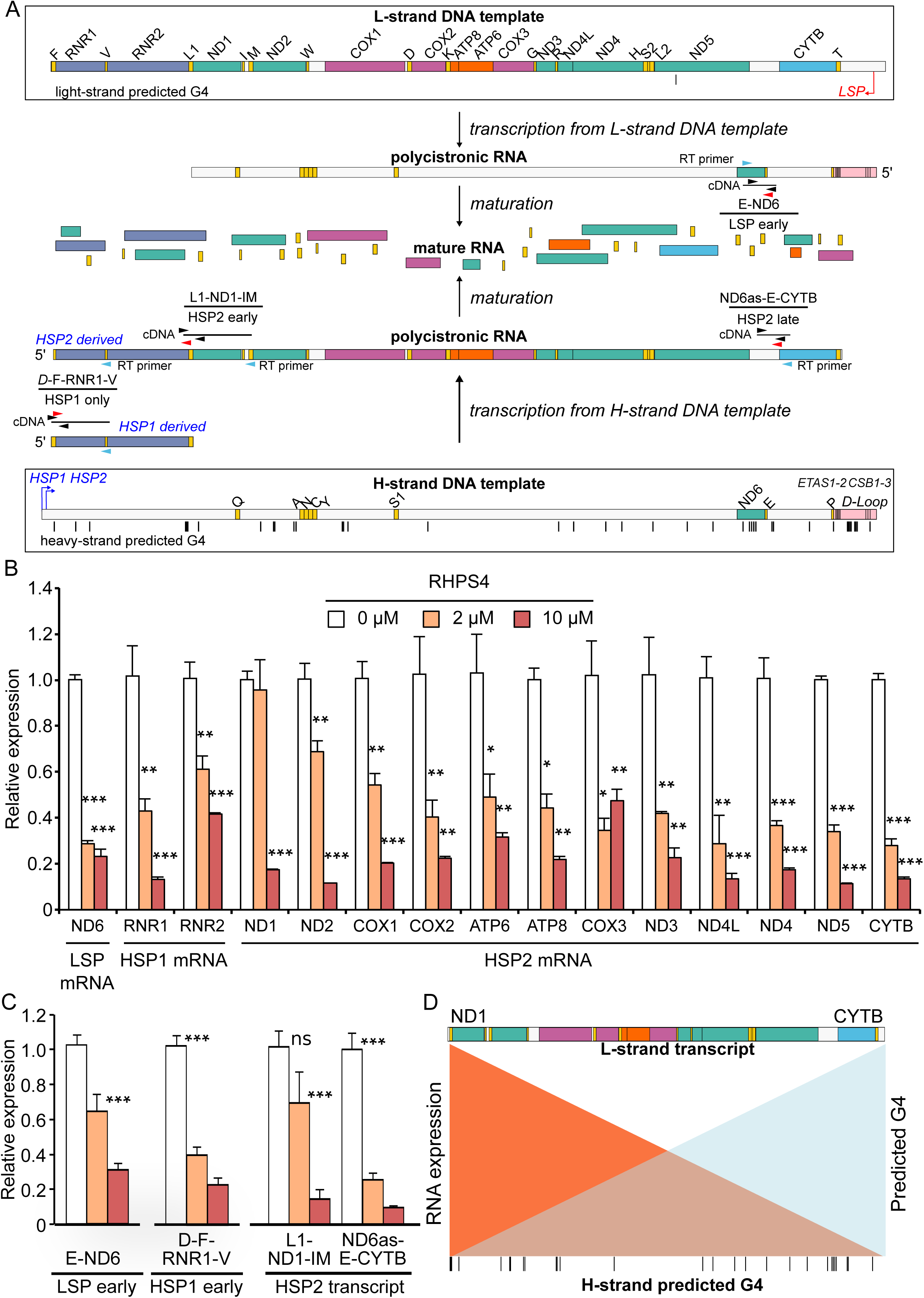
MEF exposure to low-dose RHPS4 causes transcript elongation defects prior to significant mtDNA depletion. (A) Schematic representation of mouse mitochondrial polycistronic transcripts and novel specific qRT-PCR assays. Transcription and processing of RNA proceeds from top (H-strand DNA template) or bottom (L-strand DNA template) of panel toward middle as indicated by black arrows. Below the genes are the indicated promoters (Light Strand Promoter, LSP [in red]; Heavy Strand Promoter 1, HSP1 [in blue]; Heavy Strand Promoter 2, HSP2 [in blue]). Transcription from the HSP2 promoter reads the H-strand template DNA (G-rich) to generate L-strand polycistronic RNA (G-poor), proceeding from left to right as a single transcript that is rapidly processed into mRNAs. (B) RHPS4 exposure decreases steady-state mature RNA levels assessed by qRT-PCR more extensively with distance from the promoter. Each transcript is normalized to GAPDH levels, which were unaffected by RHPS4 exposure. (C) HSP2-derived unprocessed transcript assays L-ND1-M and ND6as-E-CYTB confirm elongation defect in the presence of RHPS4. In contrast, LSP early and HSP1 early transcript assays, E-ND6 and D-F-RNR1-V respectively, show potential early transcription interference. (D) Model of progressive interference in HSP2-derived unprocessed transcript. All data are mean +/- SEM (n=3-6; p-values calculated by one-way ANOVA: *<0.05, **<0.01, ***<0.001). Mouse mtDNA sequence was used to predict quadruplex forming potential (QFP) using G4 Hunter (29).

Given the decrease in abundance of transcripts with distance from the HSP2 promoter, we hypothesized that G4s might be interfering with the elongation of transcripts by RNA polymerase. To examine elongation defects in unprocessed mitochondrial RNAs, new assays were required. Standard TAQMAN assays are intragenic, detecting both mature and nascent unprocessed RNAs, with unprocessed transcripts typically contributing a tiny portion of the total mitochondrial RNA (57, 58). To detect the nascent RNA levels, we developed strand-specific intergenic TAQMAN assays that measure unprocessed RNAs only (Figure 6A). Assays for LSP early (E-ND6), HSP1 only (D-F-RNR1-V), and HSP2 early (L1-ND1-IM) showed similar abundance at 2 μM RHPS4 (Figure 6C) to corresponding mature RNAs (Figure 6B). As with the mature RNAs, we observed reduced abundance of HSP2 late unprocessed RNA (ND6-as-E-CYTB) relative to HSP2 early sequence (Figure 6C). Because the number of potential G4-forming sequences encountered by an elongating polymerase increases with the distance from the promoter, we suggest that the observed HSP2 transcription elongation defect is caused by RHPS4-stabilization of G4 structures in the template strand (Figure 6D). A notable exception is the reduced abundance of ND6 and RNR1/2 mature RNA levels, which are generated by non-HSP2 promoters and appear to be affected by an unprocessed transcript alteration 5’ to those sequences.

Due to the relevance of transcription to replication priming (1), we also examined transcripts associated with replication priming in the RNA-seq data for RHPS4 effects. Notably, decreased strand-specific RNA abundance at the D-loop (the site of first strand replication initiation) and O_L_ (the site of second strand replication initiation) was detected at 10 μM, but not 2 μM, RHPS4 (Figure S9C and D), suggesting that the formation of replication-associated transcripts was not impeded at the lower levels of RHPS4 that do not cause mitochondrial genome depletion. We noted that at 10 μM RHPS4 exposure, the distribution of RNA transcripts around CSBI, a typical RNA-DNA transition site, was altered (Figure S9D), suggesting that the robust mtDNA depletion at this high level of RHPS4 could involve effects within the D-loop region. In contrast, the mild replication inhibition detected at 2 μM RHPS4 (Figure 1) apparently occurs secondary to effects other than direct action within the D-loop region.

### Mitochondrial transcription defects induced by RHPS4 are distinct from DNA intercalation effects

To exclude RHPS4-dsDNA interactions as a potential cause of the transcription defect, we treated MEF cells with ethidium bromide (EtBr), a dsDNA intercalating compound that causes mtDNA depletion (Figure 7). We identified EtBr concentrations that recapitulated the extent of mtDNA depletion observed during 10 μM RHPS4 exposure at 24 hours (Figure 7A). In contrast to RHPS4, EtBr exposure was only able to cause transcript decrease in the context of mtDNA depletion (Figure 7B). Notably, the relative abundance of mature CYTB mRNA was not lower than ND1 mRNA (Figure 7B), there were no differences in the abundance of 5’ and 3’ unprocessed HSP2 sequences (Figure 7C), and the four unprocessed RNAs from both transcripts were only decreased with mitochondrial genome depletion (Figure 7C). Thus, at the concentrations tested, EtBr and RHPS4 show distinct effects on mtDNA transcription.

**Figure 7.**
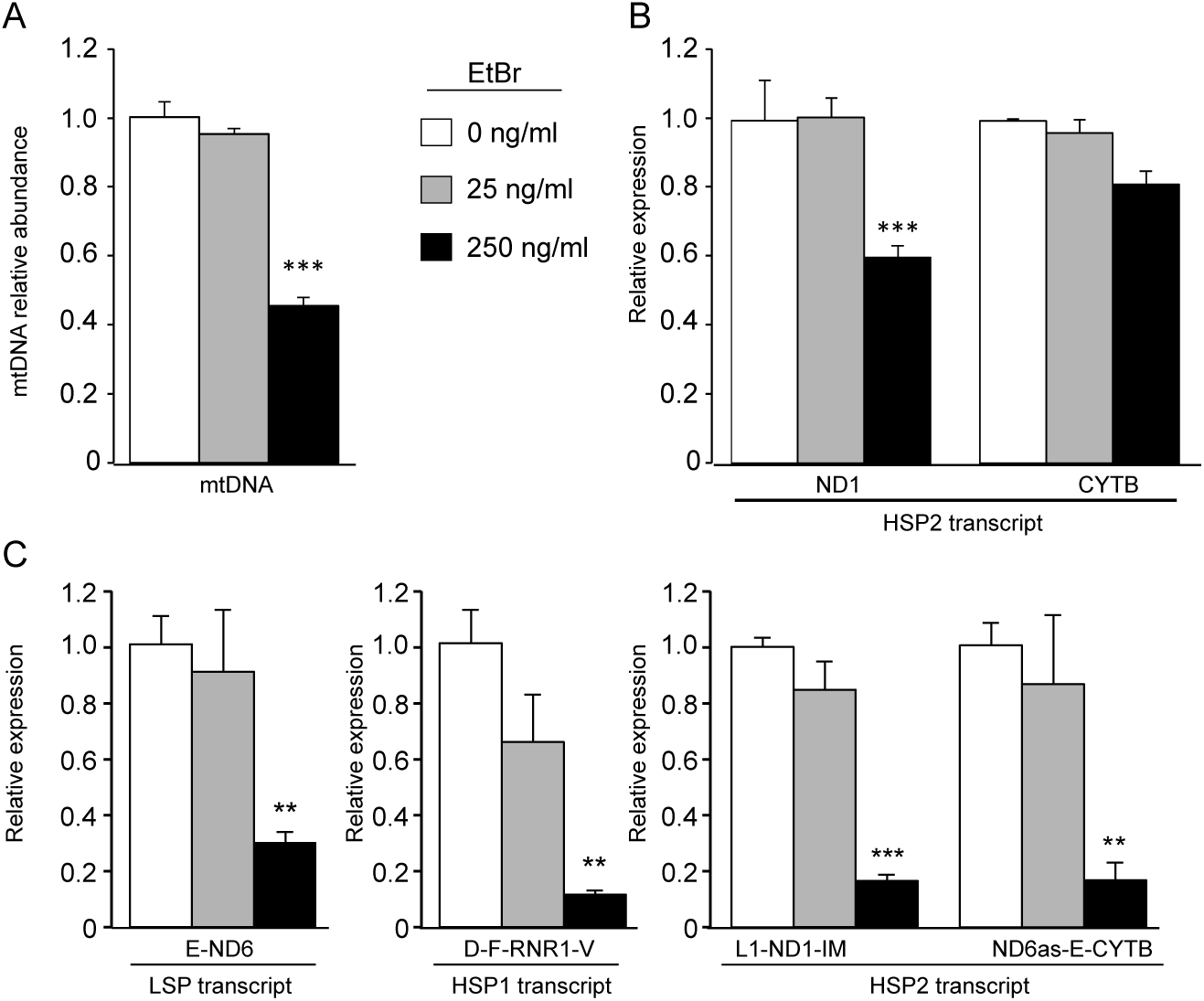
Ethidium bromide treatment does not induce mitochondrial transcription defects prior to mtDNA depletion. (A) mtDNA levels were measured in MEFs cells treated for 24 hours with 0, 25 and 250 ng/ml EtBr. Concentrations were selected to match mtDNA depletion observed in RHPS4 experiments. (B) EtBr treatment does not induce any change in the ND1 and CYTB steady-state transcript levels at 25 μM at concentrations below those that cause mitochondrial genome depletion. (C) EtBr treatment does not alter parental L-strand transcript ND6 and the H-strand transcript RNR1, ND1 and CYTB levels prior to mtDNA depletion. All data are mean normalized to untreated sample values +/- SEM (n=3; p-values calculated by one-way ANOVA: **<0.01, ***<0.001).

### Differential nuclear transcriptional responses to mitochondrial transcription inhibition and mtDNA depletion

Low and high dose RHPS4 exposure afforded the opportunity to investigate differences in cellular transcriptional responses caused by mitochondrial transcription elongation defects or mtDNA template depletion (Figure 8). Interestingly, the low and high RHPS4 treatment groups show discrete differentially expressed genes (DEG) by cluster analysis (Figure 8A). We observed that the majority of DEGs were nuclear-encoded (Figure 8B). Alterations in expression of nuclear genes encoding mitochondrial-localized proteins (based on MitoCarta2 (59)) occurred mainly at 10 μM RHPS4. We suggest that the DEGs may be largely secondary to RHPS4-mediated changes in the mitochondria because only 1 (PDGFRB) out of 9 nuclear-encoded genes with qualified G4-structures in the promoter sequences (60) showed altered expression. Enrichment analysis of the DEGs in KEGG pathways showed specific pathway transitions from low to high dose of RHPS4 (Figure 8C). The significant pathway alterations support that the DEGs are a response to mitochondrial changes. Of potential relevance to mitochondrial disorders, genes in the oxidative phosphorylation, glycolysis/gluconeogenesis, cytosolic DNA sensing/Toll-like receptor signaling, and p53 signaling pathways are altered in both RHPS4 conditions. AMPK and HIF1 signaling, steroid biosynthesis, and fatty acid metabolism, among others, showed significant pathway alterations at high dose of RHPS4. Importantly, we detected decreased gene expression affecting serine/glycine, folate, and one-carbon metabolism at 10 μM RHPS4 exposure and concomitant mtDNA depletion, which have been demonstrated to be altered in response to mtDNA replication disorders (61, 62). In contrast, genes in the mitochondrial stress FoxO signaling pathway (63), including the BNIP3 mitophagy factor (64, 65), were significantly dysregulated only at the low RHPS4 dose. Although we cannot rule out nuclear DNA damage contributing to gene expression differences, γH2AX foci formation, growth arrest, and telomere dysfunction were stimulated by RHPS4 exposure in cancer cell lines, but not in normal fibroblasts (66). The molecular determinants of the differential pathway activation between low and high RHPS4 exposure are unknown at this time.

**Figure 8.**
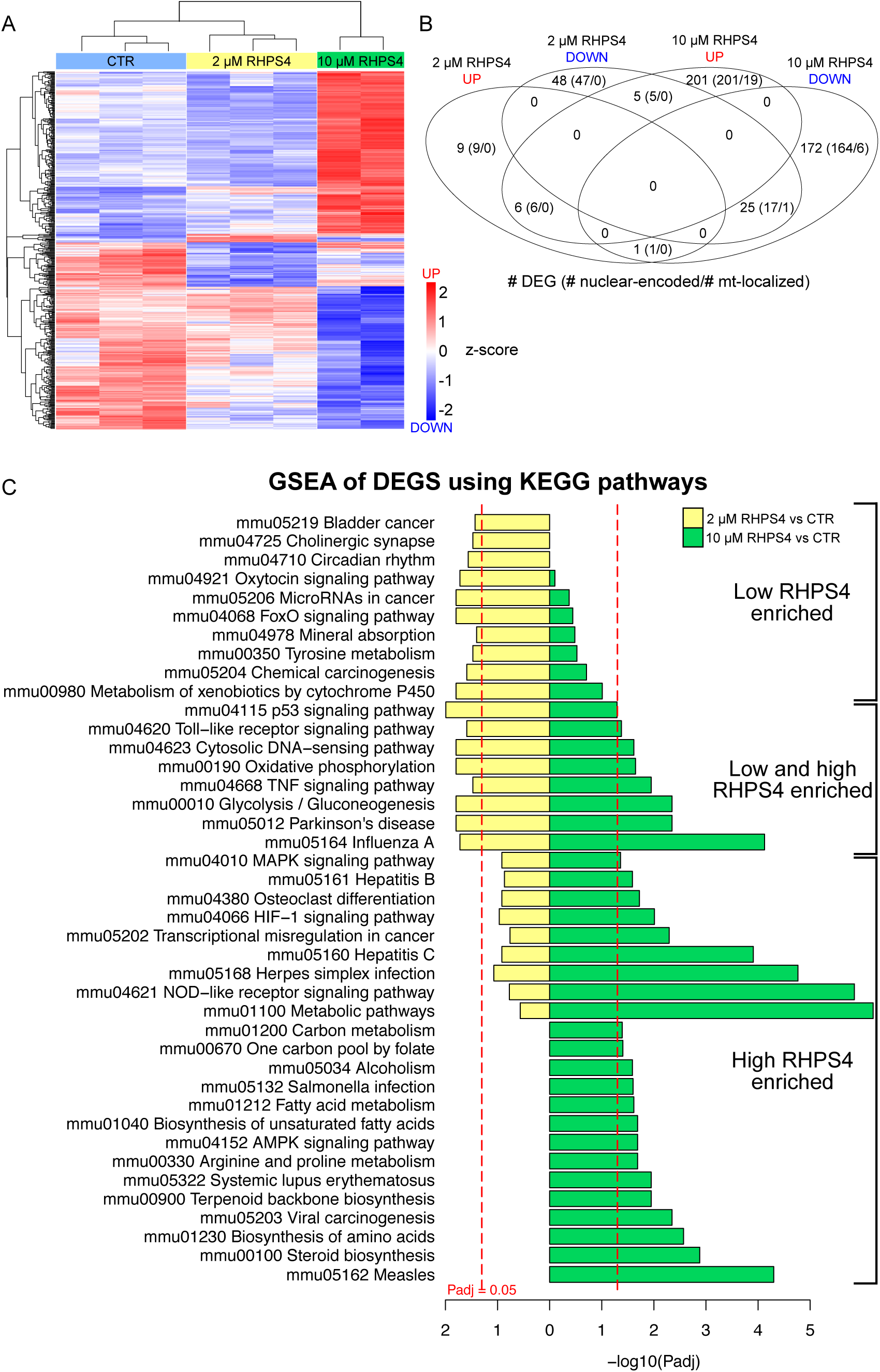
Low and high dose RHPS4 reveals differences in cellular response to mitochondrial transcription inhibition and mtDNA depletion. (A) Cluster analysis of differential expressed genes (DEGs) determined by RNA-SEQ shows discrete patterns among control (light blue), 2 μM RHPS4 (yellow), and 10 μM RHPS4 (green) treated cells. (B) Venn diagram of the number of DEG up regulated (red) or down regulated (blue) in low and high RHPS4 exposure. Subset of DEGs that are nuclear encoded genes and further subset of genes whose products are localized in mitochondria (based on the Mitocarta 2.0 database) are shown in parenthesis. (C) Enrichment of DEGs using KEGG pathways. Pathways are separated by low RHPS4 specific, low and high RHPS4 activated, and high RHPS specific changes.

### Sequence variation that increases G4 formation potential increases RHPS4-mediated respiratory defects

We tested whether mtDNA sequence could influence RHPS4-mediated effects on mitochondrial function (Figure 9). To that end, we identified the mtDNA mutation m.10191T>C, which extends one of the G-rich stretches in the predicted G4-forming sequence in the heavy strand by another guanine. The proximal region contains a G4 predicted by G4 Hunter that folds into a stable G4 structure (29) and is present in the ND3 amplicon shown to be sensitive to RHPS4 in the PCR stop assay (Figure 2E). We found that the m.10191T>C mutation increased antiparallel character of G4-structure and its stability by 8.2 °C (Figure 9A and B). To test whether RHPS4 preferentially affects this sequence, we cultured patient fibroblasts harboring the m.10191T>C mutation, or the m.13513G>A mutation (which is not expected to alter G4 structures) or control fibroblasts, with or without 1 μM RHPS4 overnight and measured oxygen consumption (Figure 9C and D). Untreated cells did not show significant differences in basal or maximal respiration among the different cell lines (Figure 9C and D). However, low dose RHPS4 treatment caused specific and profound decreases in oxygen consumption in the m.10191T>C mutation fibroblasts with no effects observed in either control or m.13513G>A mutation cell lines. These data show the potential susceptibility of specific mtDNA sequences to form G4 structures after RHPS4 treatment and regulate mitochondrial function.

**Figure 9.**
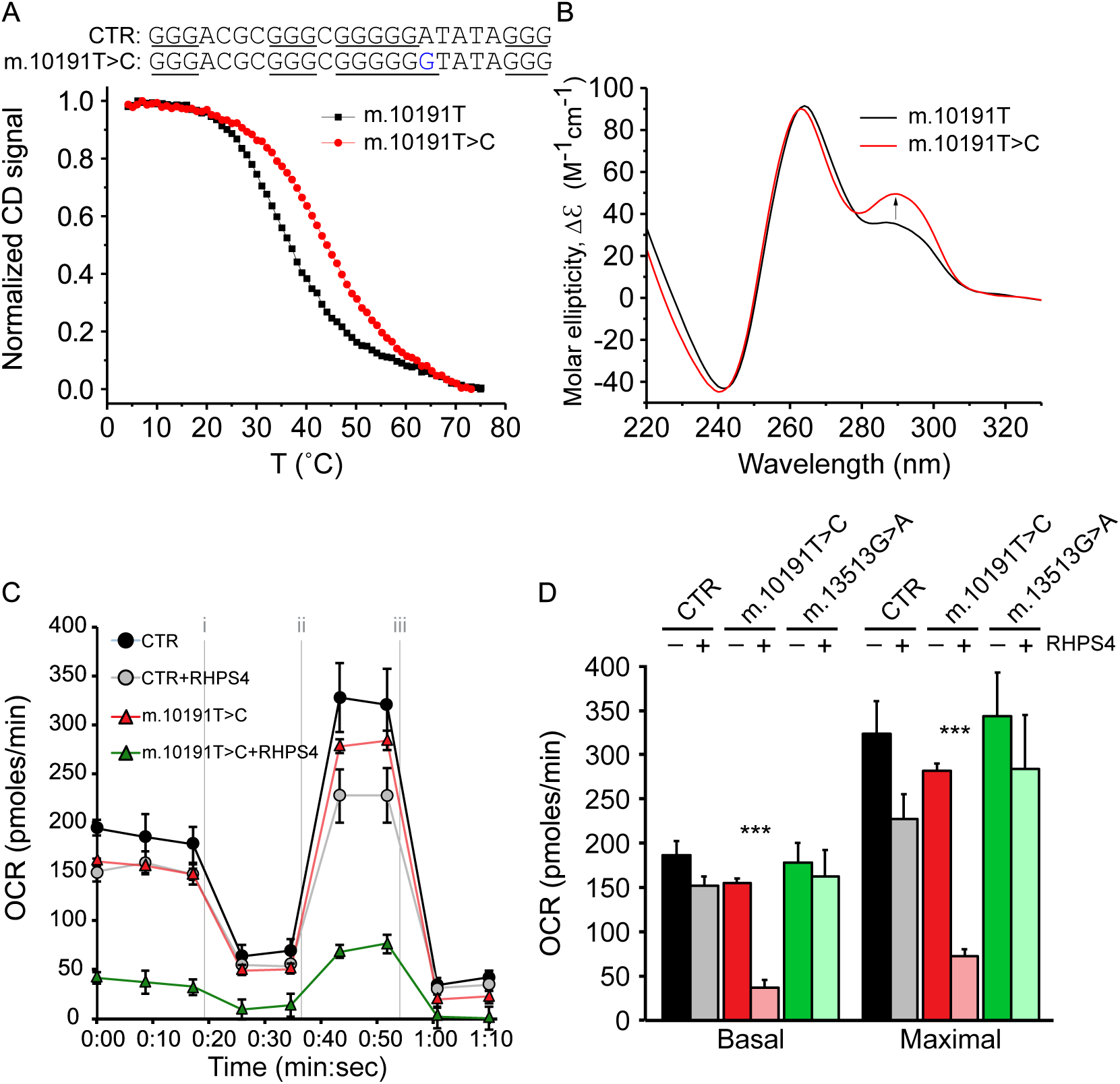
Cells harboring a mtDNA mutation that increases the G4-forming potential show oxygen consumption defects upon RHPS4 exposure. (A) CD melting curves for control (CTR) oligo nucleotide sequence m.10184-10207 and m.10191T>C mutation from heavy strand mtDNA. (B) CD spectra for CTR or the m.10191T>C mutation show differences in parallel and antiparallel character. (C) Example Seahorse Xf24 oxygen consumption rate profile for control and m.10191T>C cell lines cultivated for 48 h in presence or absence of 1 μM RHPS4 (n=5 well/line/RHPS4 condition). Treatments were as follows: i) oligomycin; ii) dinitrophenol; and iii) rotenone. (D) Effects of RHPS4 exposure on basal and maximal oxygen consumption relative to control. Included are results from cells carrying the m.13513G>A mutation, whose sequence change is not expected to alter G4 formation. All bar graph data are mean +/- SEM (p-values calculated against the untreated equivalent by Student’s two-tailed t-test: ***<0.001).

### Discussion

In this study, RHPS4 serves as an important tool for studying the effect of stabilized mitochondrial G-quadruplexes. Based on the data present here, we propose that RHPS4 translocates preferentially to mitochondria through a membrane potential-dependent mechanism, where it binds and is retained by mitochondrial G-quadruplex structures to interfere with transcription, replication, and ultimately mitochondrial respiration. The differences between our observations of mitochondrial RHPS4 localization and previous reports are well-explained in our supporting data.

Although the specific sequences bound by RHPS4 in living cells are not known, clues arise from the data presented here. Our evidence demonstrates that RHPS4 can preferentially interfere with polymerase amplification of specific regions of the mitochondrial genome, notably ND3. This region contains a predicted G4 that we demonstrated can fold into a *bona fide* G4 structure *in vitro*. Importantly, sequence changes that increase the thermal stability and antiparallel characteristics of this G4 in ND3 enhance sensitivity of cells to RHPS4, including greatly increasing the subsequent mitochondrial respiratory defect, which provide strong evidence that RHPS4 is a mtDNA structure-specific drug.

Additional sequences are likely to be involved in the RHPS4 effects on mitochondria. Broadly speaking, G4 structures can form in single-stranded nucleic acids, which form naturally during first strand replication of mtDNA. The HSP2-derived transcript elongation defects caused by low-dose RHPS4 coincide with G4-predicted sequences residing almost exclusively in the heavy strand template. We predict that the compound binds to the displaced heavy strand, which occurs in in the major arc region between the heavy and light strand origins (O_H_ and O_L_, respectively) containing two-thirds of the genome during first strand replication (26). This same region acts as the template for HSP2-derived transcription, which is also affected by RHPS4. This region also contains the majority of mtDNA deletion breakpoints, which are associated with G4-forming sequences (6) and targeted single-strand breaks in a G4-forming sequence downstream of O_L_ are sufficient to cause mtDNA deletions (22). The spontaneous formation of G4 at that sequence may similarly arrest replication and increase deletion formation. Further studies are required to reliably detect G4 structures *in situ* and identify high-efficiency forming sequences in normal and variant mtDNA sequences. Taken together, our current data suggest that conserved mtDNA strand asymmetry and consequent G4 sequence distribution can play a role in replication and gene expression.

The RNA priming of first strand mtDNA replication is thought to occur through mitochondrial RNA polymerase POLRMT-based LSP synthesis, where the nascent RNA can either elongate to generate full transcripts or terminate at the three conserved sequence blocks (CSB I, II, and III) that have been described *in vitro* (67) to provide a 3’-OH for initiation of DNA replication. Recently, transcription termination at the human CSBII has been shown to be dependent on G4 structure formation (23, 24), and prevented by the transcription factor TEFM (68, 69). As such, it might be expected that G4 stabilization would lead to transcription arrest at CSBII and LSP transcript depletion. Indeed, we observed a decrease in mature and unprocessed ND6 LSP transcripts at the low RHPS4 dose. Furthermore, the absence of RHPS4 effects at the mouse CSBII region may be due to the smaller central guanine run, consistent with the observation that the shorter central run in humans causes RNA pausing less frequently than the longer human variant (68). Further investigations and the development of site-directed mtDNA mutagenesis technologies may be required to test the function of individual G4 sequences in the replication switch.

In our working model, formation of G4 structures may be a stochastic process, but their stability mandates dedicated activities for their resolution. Potentially significant regulators of G4 structure stability are G4-resolving helicases. Among the most efficient G4 helicases is PIF1 (18). PIF1 localization occurs through alternative translation initiation mechanisms in both yeast and mammals (21, 70). PIF1 G4-helicase activity is conserved from bacteria to mammals, and is essential for mtDNA stability in yeast (71). Interestingly, skeletal muscle tissue isolated from PIF1 knockout mice shows precocious mtDNA deletions and decreased enzymatic activity in complex I (20), whose subunits are encoded by 44% of the mitochondrial genome sequence. We do not yet understand the mechanisms underlying these phenotypes and the role of G4 structures in these processes.

Although the association of G4-forming sequences with mtDNA deletion breakpoints has been demonstrated (6–8), the natural function of these structures in mitochondrial biology remains elusive (26). Using a specific G4 ligand that stabilizes G4 structures in mitochondria, we have begun to understand the impact of these sequences on mitochondrial function. While our approach likely increases both the number and stability of G4 structures, other *in vivo* events could modulate G4 formation, thereby enabling G4 dynamics to act as a mitochondrial regulatory mechanism. Such modulators could involve ATP-dependent helicases (such as PIF1 (20)), oxidative modification (8-oxoguanine and hydantoin (72, 73)), or mtDNA mutation (this study). This work will enable future studies to identify, quantify, and eventually target G4-forming sequences to increase our understanding of processes that regulate mtDNA transcription and replication as part of mitochondrial biogenesis and function.

## DATA AVAILABILITY

RNA-SEQ data are available from GEO using the token GSE103116.

## Supplementary Data

Supplementary data Figures S1-S8 are available at NAR online.

## ACKNOWLEDGEMENTS

We would like to thank Steven Barrett for some preliminary biophysical work on this project.

## AUTHOR CONTRIBUTIONS

BAK, FBJ and LAY designed experiments and reviewed data. MF, JEK, LS, CW and YVT performed cell culture experiments under BAK supervision. JT and IMX performed biophysical analysis under LAY supervision. NS and JEK performed Seahorse measurements. MF, CW and CMstC performed live cell microscopy. WH performed RNA-SEQ and TW analyzed resultant data sets.

